# Mpox mRNA-1769 Vaccine Inhibits Orthopoxvirus Replication at Intranasal, Intrarectal, and Cutaneous Sites of Inoculation

**DOI:** 10.1101/2024.09.19.613928

**Authors:** Catherine A. Cotter, Maxinne A. Ignacio, Jeffrey L. Americo, Patricia L. Earl, Eric M. Mucker, Jay W. Hooper, Tiffany R. Frey, Andrea Carfi, Alec W. Freyn, Bernard Moss

## Abstract

The increasing incidence of mpox in Africa and the recent global outbreak with evidence of sexual transmission have stimulated interest in new vaccines and therapeutics. Our previous study demonstrated that mice immunized twice with a quadrivalent lipid nanoparticle vaccine comprising four monkeypox virus mRNAs raised neutralizing antibodies and antigen-specific T cells and were protected against a lethal intranasal challenge with vaccinia virus. Here we extended these findings by using live animal imaging to demonstrate that the mRNA vaccine greatly reduced virus replication and spread from an intranasal site of inoculation and prevented detectable replication at intrarectal and cutaneous inoculation sites. Moreover, considerable protection was achieved with a single vaccination and a booster vaccination enhanced protection for at least 4 months. Protection was related to the amount of mRNA inoculated, which correlated with neutralizing antibody levels. The role of antibody in protection was demonstrated by passive transfer of immune serum pre- or post-challenge to immunocompetent and immunodeficient mice lacking mature B and T cells and therefore unable to mount an adaptive response. These findings provide insights into the mechanism and extent of mRNA vaccine induced protection of orthopoxviruses and support clinical testing.

## Introduction

The increased incidence of mpox in Africa and the recent global outbreak highlight the need for effective vaccines and therapeutics. Monkeypox virus (MPXV) and variola virus (VARV), which are responsible for human smallpox and mpox, respectively, belong to the orthopoxvirus genus within the chordopoxvirus subfamily of the Poxviridae ^1^. The conservation of essential genes among orthopoxviruses contributes to the efficacy of smallpox vaccines derived from vaccinia virus (VACV), the prototype orthopoxvirus ^2^. Jynneos, consisting of the replication defective modified VACV Ankara (MVA), was approved for mpox and smallpox prophylaxis based on animal studies as well as immunogenicity and safety in clinical trials ^3^. During the recent global outbreak of mpox, one and two vaccinations with Jynneos provided 36% to 75% and 66% to 89% effectiveness in preventing mpox, respectively ^4–6^, though anti-MPXV neutralizing antibody was low ^7^. Lipid nanoparticle (LNP) vaccines containing mpox virus (MPXV) antigen-expressing mRNAs have recently been described ^8–12^ and higher neutralizing antibody to MPXV and VACV compared to MVA was demonstrated ^8^. Two mpox mRNA vaccines are at early stages of clinical testing (NCT05995275, NCT05988203).

The development of mRNA and subunit vaccines requires an understanding of the biology of orthopoxviruses. Two related types of infectious virus particles exist: the stable mature virion (MV) is thought to be responsible for spread between hosts and the enveloped virion (EV) with an additional fragile membrane enables spread within a host ^13^. The protein components of the outer membranes of the two forms of virus differ and animal studies indicate that antibodies to at least one MV membrane protein and one EV membrane protein are needed for optimal protection against infection ^14–17^. The Moderna mRNA-1769 vaccine used in the present study consists of four mRNAs: two that express the MPXV MV proteins M1 and A29 (homologs of VACV L1 and A27) and two that express the MPXV EV proteins A35 and B6 (homologs of VACV A33 and B5) ^18^. An initial proof of concept study demonstrated that the mRNA vaccine induces antibodies in mice that neutralize VACV and MPXV and prevents VACV spread in vitro, as well as stimulating antigen specific CD4+ and CD8+ T cells. Furthermore, two intramuscular (IM) immunizations fully protected mice against weight loss and death following an intranasal (IN) infection with VACV ^18^. Here we provide a more in-depth study that evaluates protection following a single immunization as well as extended protection following a booster vaccination. By using a recombinant VACV expressing firefly luciferase for live animal imaging, we show that extreme weight loss and death follow the spread of virus from the intranasal (IN) site of inoculation to the chest and abdomen of control mice. mRNA immunization greatly reduced virus replication at the site of inoculation and prevented virus spread. There is evidence that MPXV spreads by contact and has been disseminating through mucosal routes linked to sexual activities. Here, we demonstrated that intramuscular (IM) mRNA vaccination prevented VACV replication at intrarectal (IR) and cutaneous sites of inoculation. The contribution of mRNA-1769 induced antibody to protection was determined by passive inoculation of immune serum either before or after challenge. Immune serum alone conferred protection to immune-competent and immune-compromised animals and resistance to infection correlated with the neutralization titers of the serum transferred.

## Results

### Immunogenicity and protection of mice following a single vaccination

Although many vaccines are given as a prime and boost, protection after the first dose is desirable particularly during an outbreak. To investigate protection after a single dose of the quadrivalent mRNA-1769 (referred to as mRNA hereafter), we immunized mice IM once with either mRNA or MVA for comparison and bled them 3 weeks later to determine the antibody response (Fig. 1A).

**Fig. 1.**
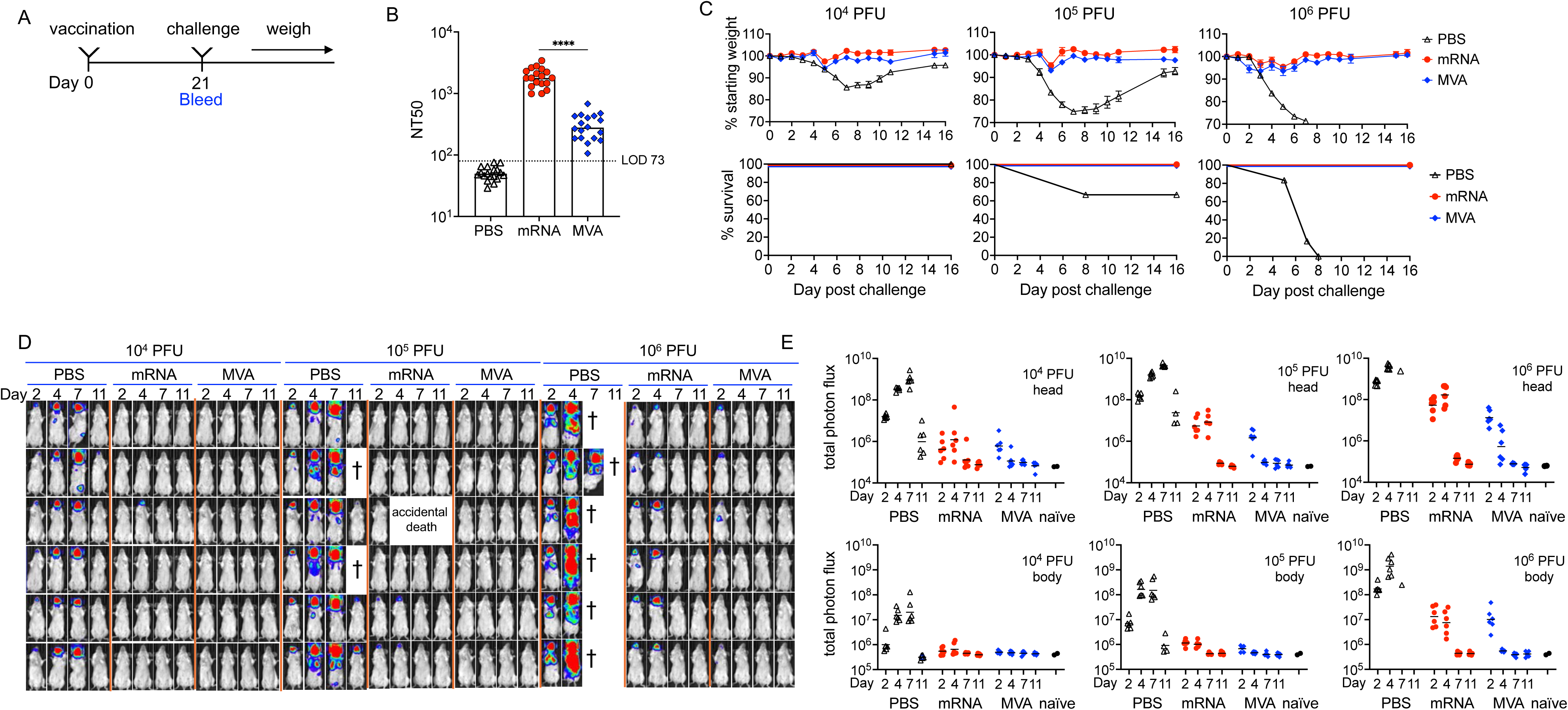
Single immunization protects against IN infection. **(A)** BALB/c mice were divided into 9 groups (n=6 per group) and mock immunized with PBS or immunized with 10^7^ PFU of MVA or 8 µg of mRNA. The mice were bled on day 0 or 21 days later prior to IN challenge with 10^4^, 10^5^ or 10^6^ PFU of WRvFire. **(B)** Anti-VACV neutralization titers expressed as NT50. **(C)** Mice were weighed daily and % of starting weight and survival plotted. **(D)** Luciferin was injected on days 2, 4, 7 and 11. BL is depicted by a pseudocolor scale with red highest and blue lowest intensity. **(E)** Photon flux was determined for head and body (chest and abdomen) ROI. The difference in peak body values between the PBS controls and the immunized mice had a significance of p≤0.0008 determined using one-way ANOVA and Tukey’s post-hoc multiple comparisons test.

Neutralizing antibodies to the M1 and A29 MV proteins are induced by the mRNAs ^8^, whereas antibodies to many proteins induced by MVA may contribute to neutralization ^19^. Nevertheless, the neutralizing antibody titer to MVs in the serum at three weeks after the mRNA immunization was a log higher than that achieved with MVA (Fig. 1B). The Western Reserve (WR) strain of VACV has been extensively used for studies of pathogenicity and a lethal dose 50 (LD50) of 10^5.3^ was determined for IN inoculation of DBA strain mice ^20^. Here we used a VACV WR recombinant (WRvFire) that expresses firefly luciferase to enable non-invasive imaging of infected BALB/c mice ^21^. The severity of infection of control BALB/c mice was proportional to virus dose leading to 30% or greater weight loss (trigger for euthanasia) or death in 0 of 6 mice at 10^4^ PFU, 2 of 6 at 10^5^ PFU, and 6 of 6 at 10^6^ PFU (Fig. 1C), resulting in a calculated LD50 of 10^5.1^. In contrast, mice receiving a single injection of mRNA or MVA vaccine exhibited minor weight loss and no deaths even at the highest challenge dose (Fig. 1C).

Bioluminescence (BL) imaging was used to determine virus replication at the site of inoculation and spread in control and immunized mice following administration of the firefly luciferase substrate luciferin on successive days following challenge (Fig. 1D). Because firefly luciferase decays with a half-life of approximately 2 h in living cells ^22^, BL reflects enzyme activity close to the time of analysis and is thus a direct measure of ongoing virus replication. In control mice challenged with 10^4^ PFU of which all survived, BL in the heads of mice reached a peak on day 7 and diminished by day 11; after challenge with 10^5^ PFU, BL was also detected in the chest by day 5 and was diminished in surviving mice by day 11; after challenge with 10^6^ PFU for which there were no survivors, BL extended to the abdomen (Fig. 1D). We concluded that weight loss is associated with upper respiratory infection and that severe disease coincides with spread to the lungs and abdominal organs in control mice. In contrast, with immunized mice BL was reduced in the head, only a few mice exhibited BL in the chest and none showed BLin the abdomen (Fig. 1D).

To avoid image saturation, a relatively short exposure time was used in Fig. 1D, and therefore could not reveal low levels of virus replication. Photon flux, however, provides a wide dynamic range and quantitative measurement of BL in regions of interest (ROI). Analysis of the head region confirmed high levels of virus replication in control mice at all challenge doses (Fig. 1E). The immunized mice exhibited a progressive increase in photon flux in the head with increasing challenge dose, but the peak values were substantially lower than in control mice and returned to baseline by day 7. Photon flux values above baseline in the bodies (which included the chest and abdomen) were detected only with the 10^6^ PFU challenge and were transient (Fig. 1E). The peak body values between the PBS controls and the immunized mice were highly significant (p<u>≤</u>0.0008) at the peak times, whereas the difference between the MVA- and mRNA-immunized mice were minimal. Thus, while neither MVA nor mRNA provided sterilizing protection at the site of inoculation when challenged with the potentially lethal 10^6^ PFU dose of VACV, replication was greatly reduced in the head and spread to the lungs and abdominal organs was largely prevented.

### Relationship of mRNA dose with neutralizing antibody and protection

Groups of mice were immunized once with 0.5 µg, 2.0 µg or 8.0 µg of mRNA. Serum was obtained at 3 weeks in each case, and also at 1 week for the 8.0 µg dose (Fig. 2A). The 3-week NT50 values were proportional to dose and the titer for the 8.0 µg dose at one week was similar to the 2.0 µg dose at 3 weeks (Fig. 2B).

**Fig. 2.**
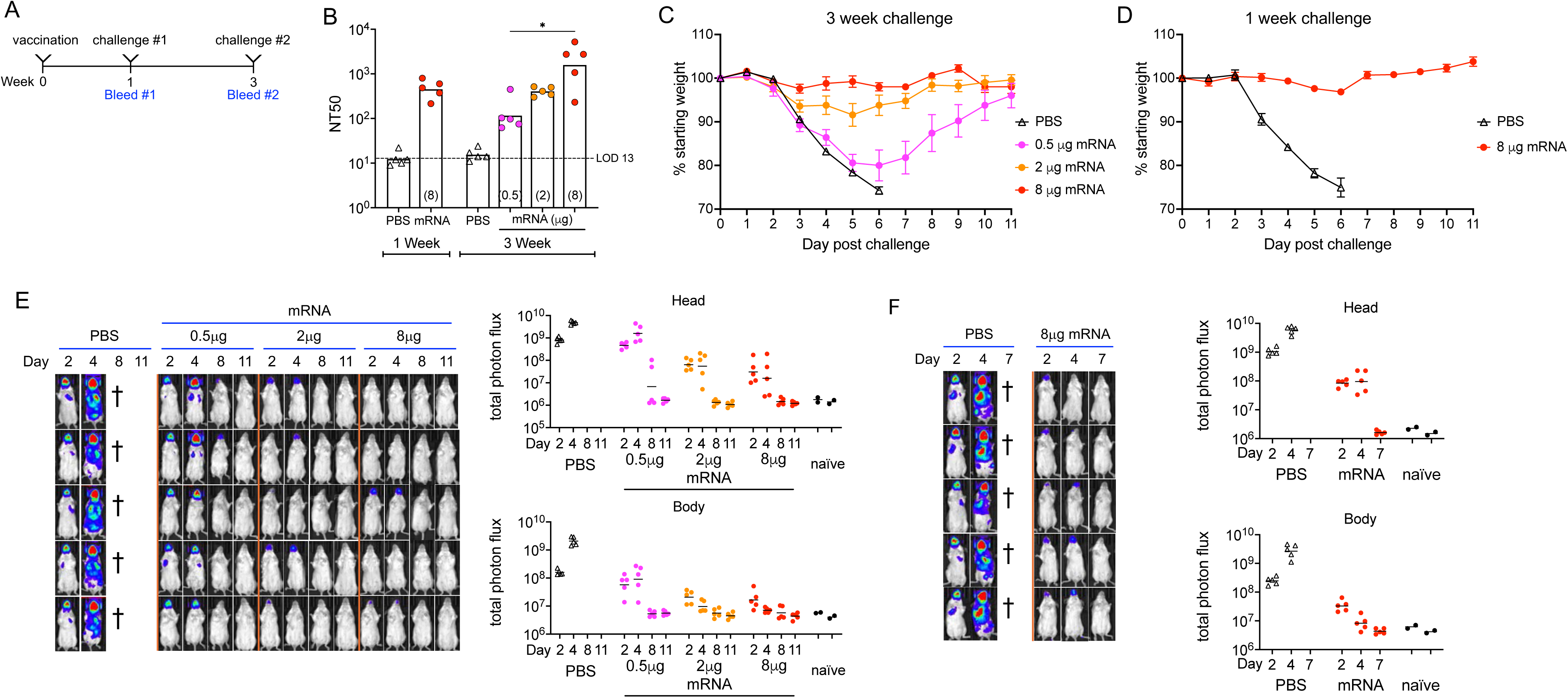
Correlation of single mRNA dose, neutralizing antibody, and protection. **(A)** Groups of BALB/c mice (n=5) were vaccinated and challenged after 1 or 3 weeks. **(B)** Anti-VACV neutralization titers expressed as NT50 determined 1 and 3 weeks after immunization with 8 µg of mRNA or 3 weeks after immunization with 0.5 and 2.0 µg of mRNA. **(C)** Mice inoculated IN with 10^6^ PFU of WRvFire at 3 weeks after vaccination with 0.5, 2.0 or 8.0 µg of mRNA were weighed daily and % of starting weight plotted. **(D)** Mice inoculated IN with 10^6^ PFU of WRvFire at 1 week after vaccination with 8 µg of mRNA were weighed daily and % of starting weight plotted. (E) Mice challenged at 3 weeks following vaccination were imaged following injection of luciferin on days 2, 4, 8 and 11. BL is depicted by a pseudocolor scale with red highest and blue lowest intensity. Photon flux was determined for head and body ROI. **(F)** Same as preceding panel except that mice vaccinated with 8 µg of mRNA and challenged after 1 week were imaged on days 2, 4, and 7.

When challenged with 10^6^ PFU of WRvFire, the control mice rapidly lost weight and succumbed to the infection, whereas the immunized mice all survived. The mice immunized with 0.5 µg of mRNA lost about 20% of their weight when challenged at 3 weeks, while the mice receiving 2 µg of mRNA lost about 6% (Fig. 2C). Mice that received 8 µg of mRNA lost no weight when challenged at 3 weeks (Fig. 2C) and lost little weight when challenged at 1 week (Fig. 2D).

The mice were imaged over a period of 11 days. As expected, the PBS control mice showed high BL in the head and extensive spread of the infection to the chest and abdomen (Fig. 2E). Mice immunized with 0.5 µg mRNA exhibited strong BL in the head and some in the chest, which was largely cleared by day 8 (Fig. 2E) consistent with the change in weight (Fig. 2C).

Mice immunized with 2 or 8 µg of mRNA exhibited low BL in the head, and none detected in the chest or abdomen (Fig. 2E). Photon flux confirmed the relatively low BL in the head and values only slightly above the baseline in the body on day 2 of mice immunized with 2 µg or 8 µg of mRNA. The BL and photon flux of mice immunized with 8 µg mRNA and challenged after one week was most similar to that of mice immunized with 2 µg mRNA and challenged at 3 weeks (Fig. 2F).

In summary, three weeks after a single vaccination, NT50 values of 10^2^ to 10^3^ achieved with 2 µg or higher mRNA resulted in little or no weight loss and reduced virus replication upon challenge with 10^6^ PFU of VACV. A similar result was achieved 1 week after a single 8 µg mRNA vaccination. Taken together these data indicated rapid protective immunity following a single mRNA vaccination that corresponded with vaccine dose and neutralizing antibody titers.

### Increased immune response and sustained protection following a boost vaccination

To determine the impact of boosting, mice received a second inoculation of 8 µg of mRNA or 10^7^ PFU of MVA at 3 weeks after the first (Fig. 3A). For each vaccine, neutralizing antibody increased after the boost and decreased slightly between 3 and 16 weeks (Fig. 3B). Again, the titers achieved with mRNA were about a log higher than with MVA.

**Fig. 3.**
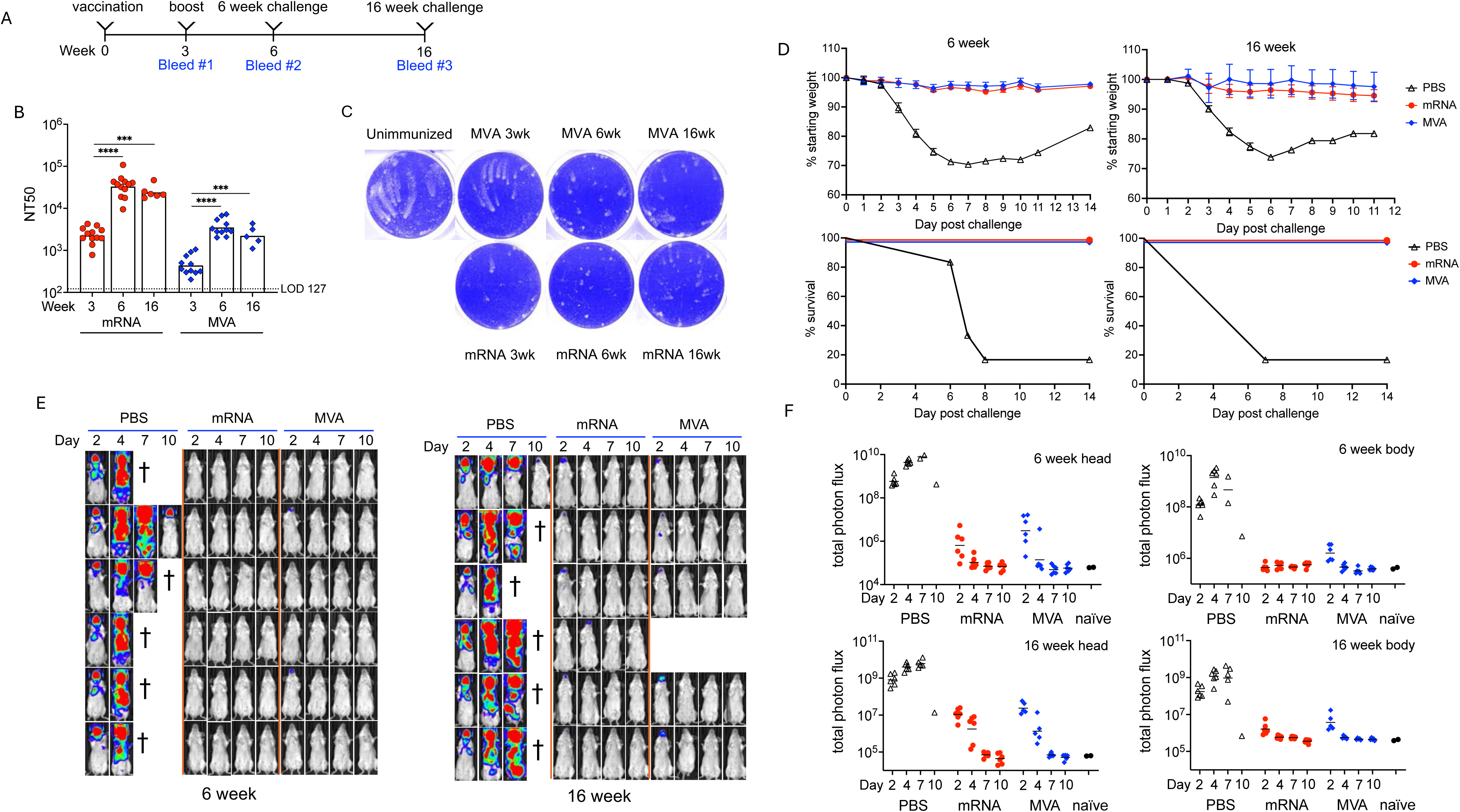
Enhanced protection after second immunization. **(A)** BALB/c mice were divided into 6 groups (n=6) and mock immunized with PBS or immunized with 10^7^ PFU of MVA or 8 µg of mRNA on days 0 and 21 and challenged on weeks 3 and 16 with WRvFire. **(B)** Anti-VACV neutralization titers expressed as NT50. **(C)** Comet spread assay. Crystal violet-stained cells shown. **(D)** % of starting weights and survival. (**E**) BLI on days 2, 4,7 and 10. (**F**) Photon flux of head and body ROI.

In this experiment, an assay to measure functional antibodies targeting the EV was conducted. Antibodies to the VACV homologs of the EV proteins A35 and B6 reduce virus spread on a cell monolayer when antibody is added after virus adsorption and the plates are overlaid with liquid medium ^16^. After incubation for 2 days, crystal violet staining revealed satellite plaques with a characteristic comet-like distribution in the presence of non-immune serum, but which were reduced partially by 3-week serum and more completely by 6-week serum (Fig. 3C). The comet inhibition decreased slightly in the serum obtained at 16 weeks (Fig. 3C).

The PBS-inoculated mice lost weight and 5 of the 6 succumbed when challenged with 10^6^ PFU of WRvFire, whereas mRNA- and MVA-immunized mice lost little or no weight and survived when challenged at 6 or 16 weeks (Fig. 3D). Although the MVA vaccine induced lower neutralizing antibodies than mRNA, it likely exceeded the threshold necessary for protection of mice.

Images made after the 6-week challenge revealed transient BL in the heads of 2 of the 6 mice that received MVA and none that received mRNA (Fig. 3E). BL was not detected in the bodies of immunized mice at week 6 (Fig. 3E). The mean photon flux levels in the heads of immunized mice were 3 and 4 logs lower than controls in MVA- and mRNA-immunized mice, respectively (Fig. 3F). The mean photon flux in the bodies of the MVA-immunized were barely above baseline and were at baseline for the mRNA-immunized mice.

When challenged at 16 weeks, BL was transiently detected in the chest of one MVA-immunized mouse and a few in each group had transient BL in the head (Fig. 3E). The mean photon flux of the heads was higher than at 6 weeks but still nearly 3 logs lower than controls and the photon flux in the bodies was only slightly above baseline on day 2 before returning to baseline (Fig. 3F). Thus, a high degree of protection was sustained for at least 4 months.

### Protection against intrarectal (IR) and percutaneous infections

Although MPXV can spread through the respiratory route, human-to-human transmission occurred mostly through male-to-male sexual activity during the recent global outbreak, and lesions in the rectal mucosa were common ^23^. Furthermore, there is new evidence of sexual spread of the more pathogenic clade I MPXV in Africa ^24^. It was important therefore to determine whether mRNA administered IM would protect against IR and percutaneous challenges. Mice were primed and boosted with mRNA or MVA vaccines and then inoculated with 10^6^ PFU of WRvFire IR. In control mice, local BL was detected on day 2, increased on day 4 with detectable BL in the abdominal region of one mouse, and was diminished by day 8 (Fig. 4A). In contrast, no virus replication was detected by either BL (Fig. 4A) or more sensitive photon flux measurements (Fig. 4B) in immunized mice.

**Fig. 4.**
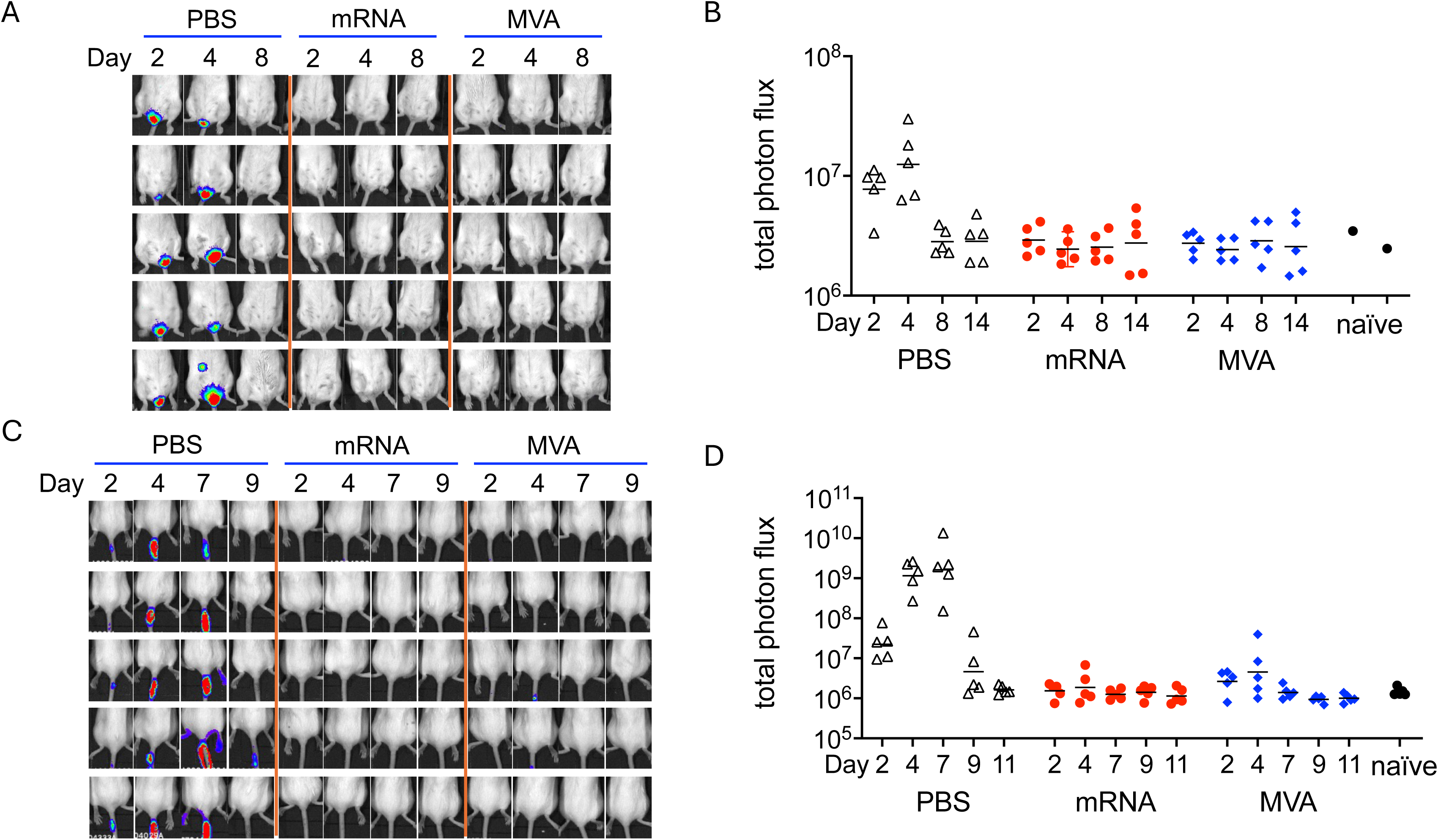
Protection from IR and percutaneous infection, **(A)** BLI of mice (n=5 per group) following IR infection with 10^6^ PFU of WRvFire. (B) Total photon flux of the rectal area of interest. (C) BLI of mice (n=5 per group) following percutaneous infection with 10^5^ PFU of WRvFire. (D) Total photon flux of the area around the tail.

The smallpox vaccine was administered percutaneously in humans and caused a pustular lesion within 7 to 8 days. Here we used a similar inoculation strategy with mice except that the pathogenic WRvFire was used instead of the attenuated vaccine strain. Following percutaneous inoculation of control mice with 10^5^ PFU of WRvFire, BL peaked on days 4 and 7 at the site of inoculation and substantially declined by day 11 (Fig. 4C, D). In contrast no BLI or increased photon flux was detected in immunized mice. Thus, IM inoculation with either mRNA or MVA protects against IN, IR and percutaneous infections in VACV mouse models.

### Passive immunization protects against subsequent virus infection

Previous studies indicated that antibody has a dominant role in protection mediated by live VACV vaccines ^25–27^. It was of interest, therefore, to determine whether mice would be protected from VACV challenge by passive transfer of serum from mRNA-immunized animals. Because only small amounts of mouse serum were available, we used serum from cynomolgus macaques that was colleted four weeks after priming and boosting with mRNA (Fig. 5A). In a preliminary experiment we determined that the titer of neutralizing antibody in the blood of mice (n=4) had a half-life of 7 days following IP inoculation. For the challenge experiment, mice were transfused with pooled serum from macaques that were immunized with 0, 15, 50 or 150 µg of mRNA (abbreviated as 0 µg,15 µg, 50 µg and 150 µg mRNA serum). The NT50 titers of the sera from mice at 1 day after transfusion were proportionate to the dose of mRNA used for immunization of the macaques, whereas the titers of the sera at 12 days were more similar (Fig. 5B) due to antibody induced by the challenge virus as will be discussed below.

**Fig. 5.**
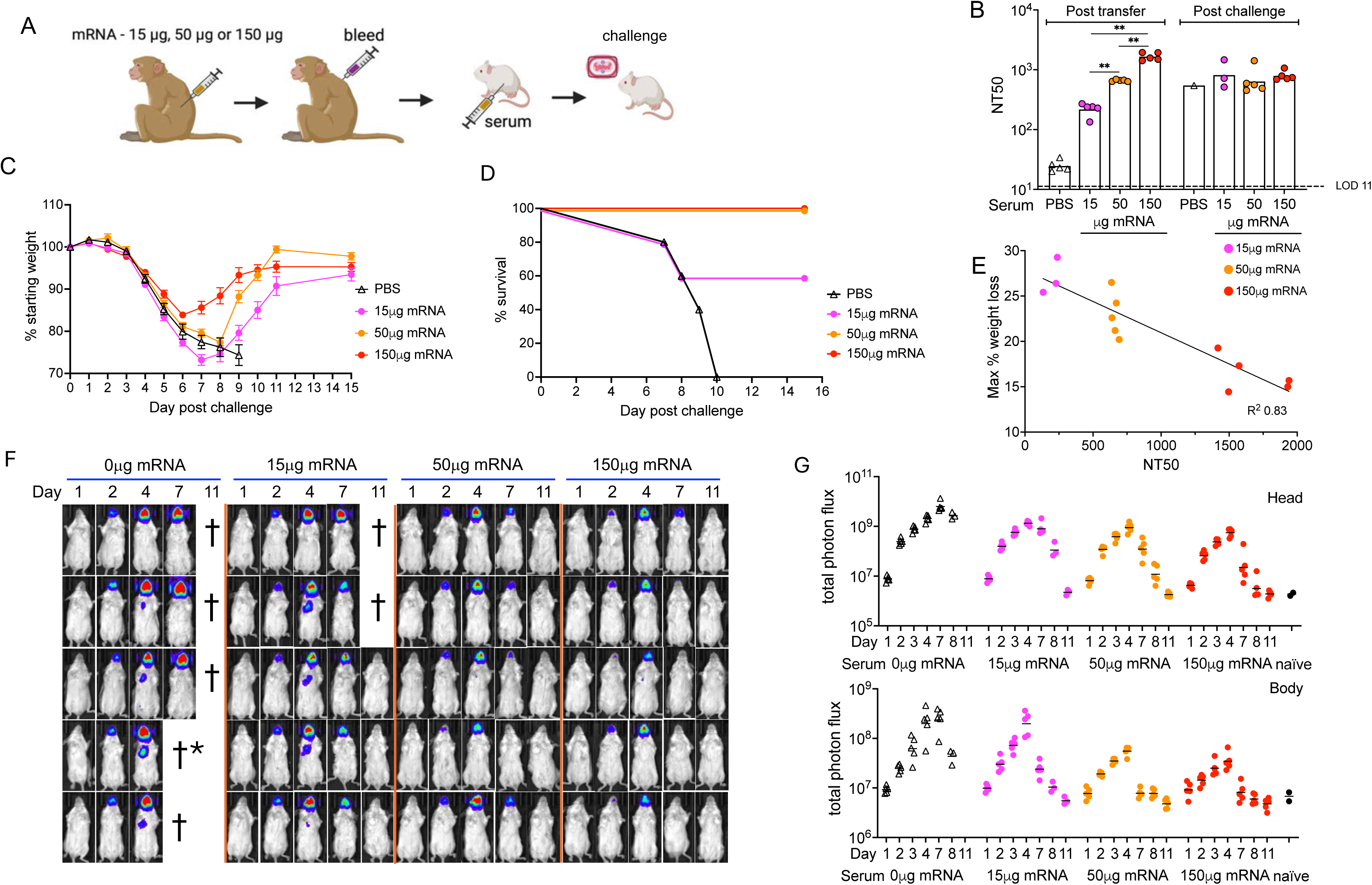
Immune serum protects against prior virus infection. **(A)** Immunization and challenge scheme. Macaques were immunized by priming and boosting with 0 (control) or 150 µg of mRNA. Mice (n=5 per group) were infected IN with 10^5^ PFU of WRvFire and pooled macaque serum was inoculated IP one day later. **(B)** Mice were weighed daily following challenge. **(C)** Mice that died naturally or euthanized because of a weight loss of 30% or more are indicated. **(D)** BLI obtained on days indicated. **(E)** Total photon flux was determined for head and body ROI.

**Fig. 5.**
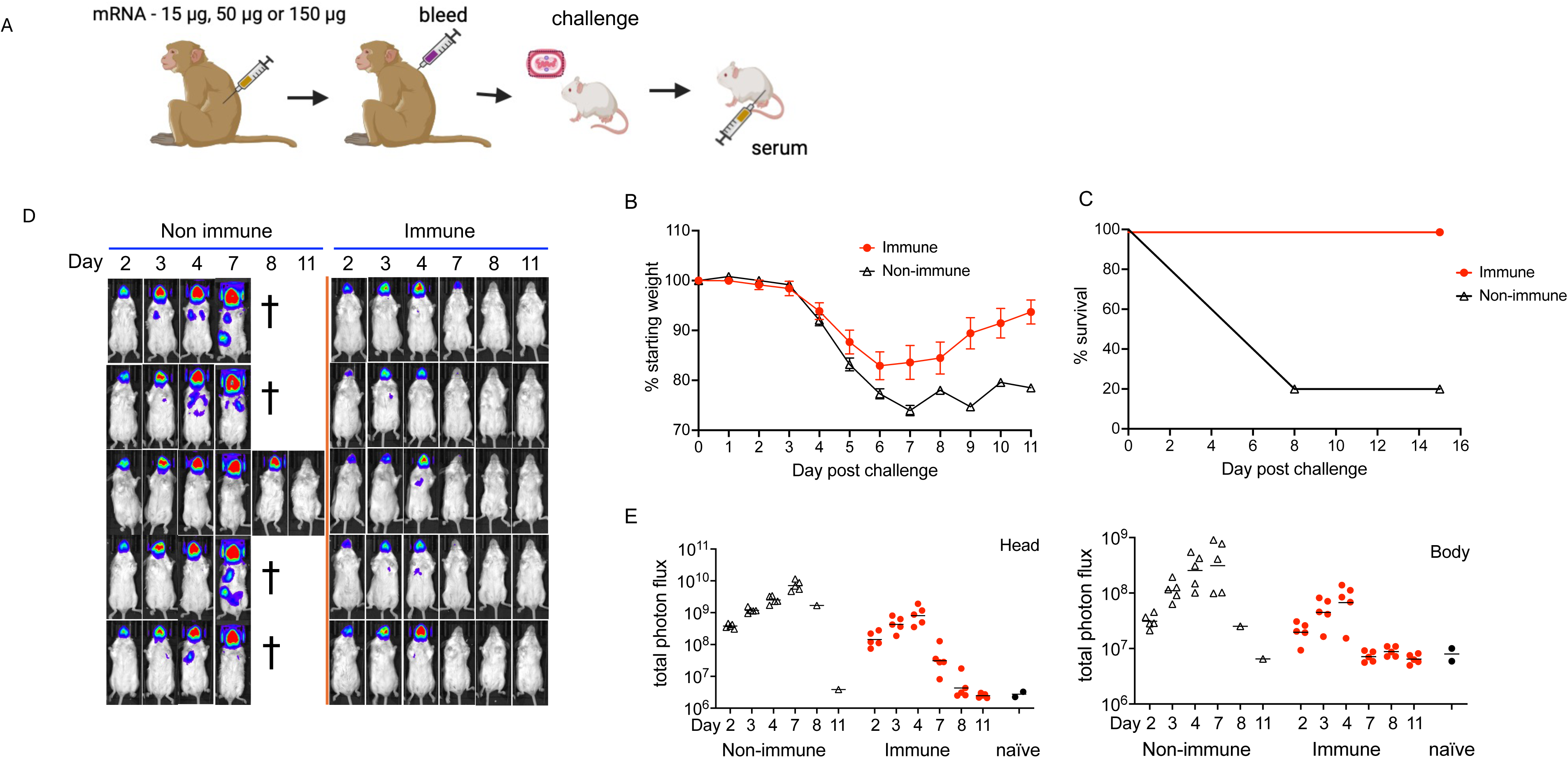
Immune serum protects against subsequent virus infection. **(A)** Immunization and challenge scheme. Macaques were immunized by priming and boosting with 0, 15, 50 or 150 µg of the mRNA vaccine. Pooled serum was injected IP into BALB/c mice (n=5 per group) and one day later the mice were challenged with 10^5^ PFU of WRvFire. **(B)** Serum was obtained from mice prior- and post-challenge and the anti-VACV neutralization titers were determined. **(C)** Mice were weighed daily following challenge. **(D)** Mice that died naturally or euthanized because of a weight loss of 30% or more are indicated. **(E)** Relationship of weight loss to NT50 is plotted. R^2^ of 0.83 was determined. **(F)** BLI obtained on days indicated. **(G)** Total photon flux was determined for head and body ROI.

The mice were challenged IN with 10^5^ PFU of WRvFire (1 log less than after active mRNA immunizations) at one day after transfusion. Mice that received control (0 µg mRNA) serum rapidly lost weight and succumbed (Fig. 5C, D). Mice that received the 15 µg mRNA serum also exhibited severe weight loss, but half recovered before losing 30% of their weight. In contrast, all mice that received sera from the macaques immunized with 50 or 150 µg mRNA sera survived, although the former suffered more weight loss before recovering. There was an inverse relationship between the neutralization titers on day 1 and maximum weight loss of surviving mice (Fig. 5E). Nevertheless, the mice receiving macaque immune serum were not as well protected against weight loss as mice that were actively immunized even though the NT50 values were in the same range at the time of challenge.

Mice that received the control (0 µg) serum exhibited intense BL in the head and chest as expected. Mice that received the 15 µg mRNA serum also showed strong BL in the head and detectable BL in the chest, although lower than in the mice that received non-immune serum (Fig. 5F). However, the mice that received the 50 and 150 µg mRNA sera exhibited much lower BL in the head and undetectable BL in the body. Photon flux measurements indicated that mice receiving the 50 and 150 µg sera had about a log less virus replication than control mice in the heads and bodies at the peak times (Fig. 5G).

Although the mice receiving control serum succumbed quickly following challenge, it was possible that an immune response to the challenge virus contributed to the protection of mice receiving immune serum. To investigate this possibility, we determined the neutralizing antibody titers of the surviving mice (Fig. 5B). Mice that received the 15 µg mRNA serum and recovered exhibited a 6-to 13-fold increase in NT50; mice that received the 50 µg mRNA serum exhibited less than a 2-fold mean increase in NT50 and mice that received the 150 µg mRNA serum had a decrease in NT50 consistent with the 7-day half-life of the macaque antibodies. Thus, an immune response to the challenge virus was unlikely to benefit mice that received the 150 µg serum, though it may have helped those that received lower titer macaque serum.

### Passive immunization provides protection post-exposure to VACV infection

The VACV infection model uses the purified MV form of the virus for challenge. However, spread of VACV within an animal depends on the EV form of VACV with different surface proteins ^28^. To determine whether the immune serum could control spread of an established infection, serum was added one day after the virus inoculation (Fig. 6A). Mice that received the 150 µg mRNA serum exhibited transient weight loss followed by recovery on day 7, whereas 4 of 5 mice that received nonimmune serum succumbed to the infection (Fig. 6B, C). The recovery of the immunized mice was also demonstrated by the diminution of BL in the head between day 4 and 7 (Fig. 6D). Photon flux measurements indicated that the peak replication in the body was about 1 log lower in mice that received immune serum compared to mice that received control serum (Fig. 6E). Taken together, passive antibody elicited by mRNA-1769 vaccination provided protection when administered before or after challenge.

**Fig. 6.**
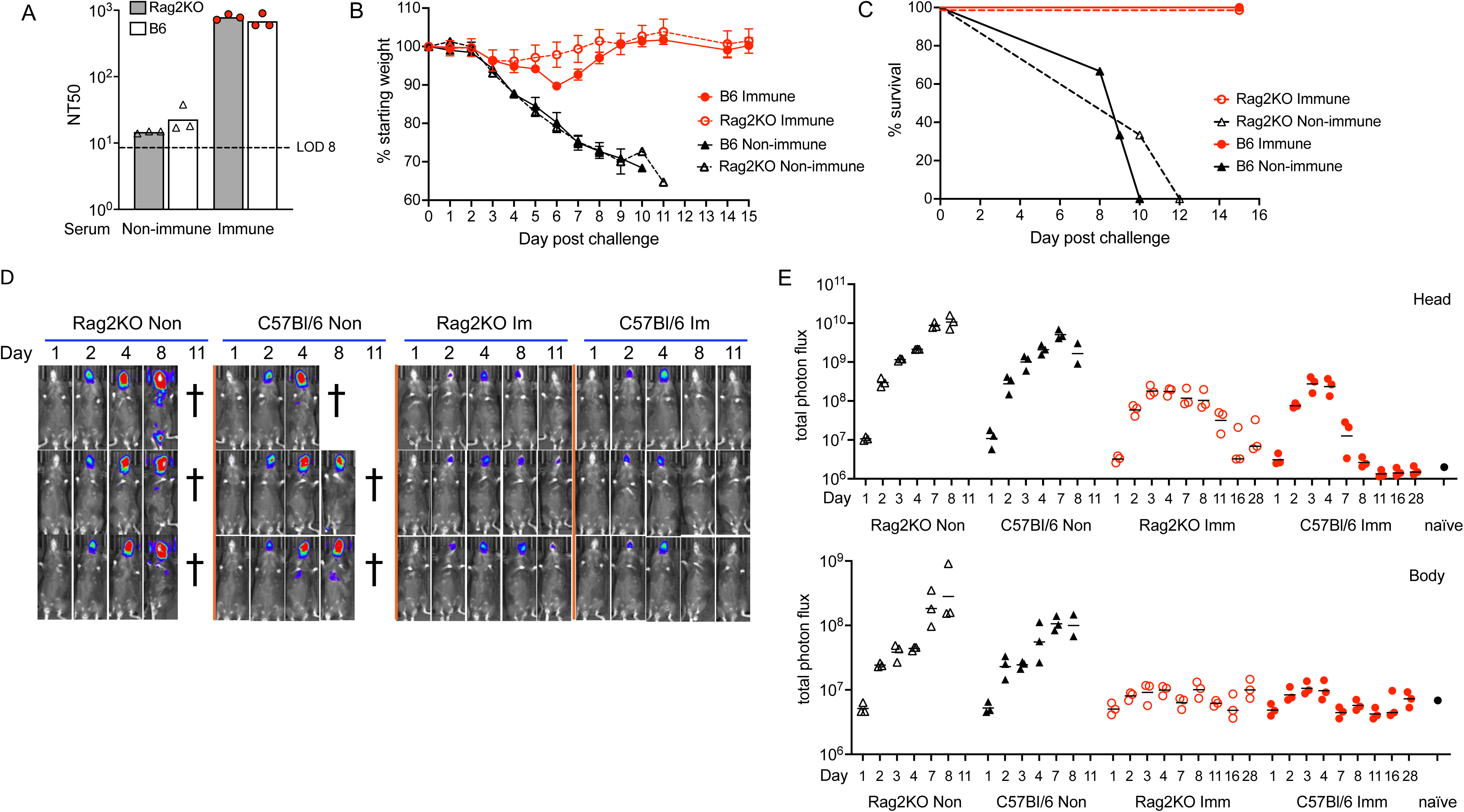
Immune serum protects immunodeficient Rag2 KO mice. **(A)** Neutralization titers of serum from mice (n=3) prior to challenge. **(B)** Mice were challenged one day after receiving control or immune serum and weighed daily. **(C)** Mice that died naturally or euthanized because of a weight loss of 30% or more are indicated. **(D)** BLI shown for days indicated. Non, non-immune; Im, Immune **(E)** Total photon flux was determined for head and body ROI.

### Passive immunization protects against acute disease in the absence of an adaptive immune response

The recovery of weight at day 7 following virus challenge (Fig. 5A) raised the possibility of a role for an immune response elicited by the infection, although protection was shown to correlate with the abundance of neutralizing antibody transferred. To investigate a role for adaptive immunity, we passively transferred control 0 µg or 150 µg mRNA immune serum to C57Bl/6 Rag2 (recombination activating gene 2) knock-out (KO) mice, which have a deletion of the entire RAG2 protein coding region and consequently produce no mature T cells or B cells ^29^. As a control, immunocompetent parental C57Bl/6 mice also received control and immune serum. The passively administered neutralizing antibody titers were similar in both mouse strains as measured one day post transfer (Fig. 7A). Following challenge with 10^5^ PFU of WRvFire, the C57Bl/6 and Rag2 KO mice that received the control serum rapidly lost weight and succumbed to the infection (Fig. 7B, C). By comparison, Rag2 KO and C57Bl/6 mice that received the immune serum lost relatively little weight and all survived until the experiment was ended at 28 days. A previous study found that BALB/c SCID mice passively immunized with rabbit polyclonal anti-L1, -A33 and -B5 antibodies had a 50% mean survival time of 26 days following IN challenge with VACV ^16^.

Live imaging provided information regarding virus replication. BL was intense in the heads of mice that received control serum and was also detected in their chests, whereas the BL was less intense in the heads and undetectable in the bodies of Rag2 KO and C57Bl/6 mice that received immune serum (Fig. 7D). Furthermore, the mean photon flux was up to 2 logs lower in the heads of mice that received immune serum compared to those that received non-immune serum and was barely above baseline in the bodies of the former (Fig. 7E). However, in the immunocompetent mice virus was cleared from the head by day 11, whereas in the Rag2 KO mice luminescence, indicating the presence of live virus, was still detected on day 28 (Fig. 7E).

The conclusions of the passive transfer experiments are that antibody is sufficient to protect immunocompetent and immunodeficient mice from acute disease but that an adaptive immune responses is important to totally clear virus.

## DISCUSSION

The increasing incidence of mpox in Africa and the recent outbreak in more than 100 countries emphasize the need for vaccines that can provide rapid protection and be produced at a global scale. Furthermore, the current demographics indicate the importance of reducing human-to-human transmission particularly by sexual contact. Our previous study ^8^ showed that a vaccine comprised of lipid nanoparticles containing four MPXV mRNAs induce neutralizing antibodies and T cells and provide protection against weight loss and death in a murine model. A feature of the present study was the use of live animal imaging to assess replication and spread of the virus from IN, IR and cutaneous sites of inoculation. The key findings are (i) virus spread from the IN site of inoculation to the chest and abdominal organs was associated with severe disease and mortality in control mice; (ii) one or two IM mRNA vaccinations greatly reduced virus replication at the IN site and prevented systemic spread; (iii) neutralizing antibody was detected on day 7, increased at 3 weeks and persisted for at least 16 weeks after vaccination; (iv) neutralization titers and protection were mRNA dose-related; (v) replication of virus was local following IR and cutaneous inoculation of control mice; (vi) virus replication was undetectable at IR and cutaneous inoculation sites of vaccinated mice; (vii) passive immune serum administered to immunocompetent mice before or after IN challenge provided complete protection against a potentially lethal infection; and (viii) RAG2 KO mice unable to make an adaptive immune response were protected by passive antibody against acute disease, though were unable to completely clear virus from the IN site of virus inoculation by 28 days.

Live BL imaging of VACV was previously used to investigate the roles of innate immunity on VACV replication ^30–32^ and protection against lethal disease by Dryvax smallpox vaccine and VACV immunoglobulin ^33^. Here, we used BL imaging to determine the effect of an mRNA vaccine on replication and spread of VACV from diverse sites of inoculation. We tested escalating amounts of mRNA and found that at 2 µg or higher, a single mRNA vaccination prevented weight loss and death following IN challenge of mice. Animal imaging revealed that vaccination reduced replication by 99% at the IN site of inoculation. Moreover, spread to the body was undetected after challenge with a 50% lethal dose and was reduced by more than 99% with 10X lethal dose. After a second vaccination protection persisted for at least 4 months.

Although MPXV can be transmitted through the respiratory route, human-to-human spread of the virulent clade I virus in the Democratic Republic of the Congo is thought to have occurred predominantly by close contact in families ^34,35^ and more recently through sexual activities ^24^. The latter was also the predominant mechanism of spread of clade IIb MPXV during the recent global outbreak ^23^. We found that following IR inoculation of control mice with virus, replication at the site of administration was detected by luminescence on day 2, peaked on day 4 and was eliminated by day 8 without significant systemic spread, analogous to the predominantly local lesions in the current mpox outbreak. In contrast, no luminescence was detected in the mRNA-immunized mice indicating little or no replication.

Control mice inoculated percutaneously developed pustular lesions and exhibited strong luminescence that peaked on day 7. However, no lesions formed in immunized mice and BL was below detection. An mRNA vaccine was also shown to reduce BL of a vaccine strain of VACV administered subcutaneously ^11^. Thus, mRNA vaccination administered IM is able to protect mice against mucosal and cutaneous infections, the predominant mode of human-to-human transmission of MPXV.

Although we previously demonstrated stimulation of antigen-specific T cells as well as antibodies following mRNA immunization, antibodies are generally thought to be most important for vaccine-mediated protection against VACV ^25–27^. Complete protection of mice against weight loss occurred with neutralizing titers of 10^2^ to 10^3^, which could be achieved within 7 days following vaccination. To investigate protection by passive transfer of antibodies, we used serum from non-human primates that had been immunized with mRNA because only low amounts of mouse serum was available. As precedent, several prior studies demonstrated protection against acute bacterial and viral infections including VACV following passive transfer of monkey or human serum to mice ^36–39^. Because mice are inoculated with the MV form of VACV and spread also depends on the EV form ^28^, we challenged the mice by the IN route either one day after administration of serum or one day before. Presumably, the anti-MV antibodies would be most effective when present prior to exposure to MV, and the anti-EV antibodies would be crucial after initiation of infection when spread is occurring. In both cases, the mice survived, and live imaging showed clearance of virus. However, there was more virus replication and greater weight loss in mice that were passively immunized compared to those actively immunized even though the neutralizing antibody titers were in the same range at the time of challenge. This difference could be due to rapid depletion of heterologous antibody. However, we determined that the half-life of the macaque neutralizing antibodies in mice was 7 days, similar to that of mouse antibodies ^40^. Another possibility was functional incompatibility of the primate antibodies in the mouse system. Other studies, however, have shown good binding of primate IgGs to mouse Fc gamma receptors ^41^ suggesting that the macaque antibodies are functional. The stimulation of innate immunity and induction of antigen-specific B and T cells by the MPXV mRNAs ^8^ likely contribute to the superiority of active versus passive immunization.

Another feature of the passive transfer experiments was progressive weight loss up to day 7 following virus challenge, followed by recovery. A similar phenomenon had been reported in mice that received rhesus macaque immune serum and challenged with ZIKA virus ^37^. One possibility is that an adaptive immune response to the challenge virus kicked in by day 7, helping to eliminate the infection. Indeed, the surviving mice that were passively immunized with low titer neutralizing serum exhibited an increase in neutralizing antibodies. However, the mice that received high titer serum had decreased neutralizing titers consistent with decay due to the 7-day half-life of the macaque antibody. To further investigate whether an adaptive immune response is required for protection of passively immunized mice, we employed RAG2 KO mice that cannot make mature T or B cells. These mice were protected from weight loss and virus spread as well as immunocompetent mice indicating that an adaptive immune response was not required to protect against severe acute disease. However, innate immune responses are unimpaired in RAG2 KO mice and natural killer (NK) cells ^42^ likely contributed to virus clearance. Nevertheless, clearance of the virus from the site of inoculation was delayed and incomplete in the Rag2 KO mice. We conclude that antibodies induced by mRNA are sufficient for survival of a lethal infection in the mouse model. However, the better protection afforded by active immunization with mRNA-1769 is likely due to the induction of antigen-specific B and T cells. In conclusion, the rapid induction of protectice antibodies by mRNA-1769 and the protection at IN, mucosal and cutraneous sites in mice support further testing of this promising vaccine.

## MATERIALS AND METHODS

### Biosafety

All work with infectious VACV and potentially infectious material derived from animals was conducted in a Biosafety 2 laboratory at the National Institute of Allergy and Infectious Diseases, National Institutes of Health in Bethesda, MD. Sample inactivation and removal was performed according to standard operating protocols approved by the local Institutional Biosafety Committee.

### Mice

Female 5- to 6-weeks old BALB/c mice were purchased from Taconic Biosciences. Rag2 KO (B6.Cg-*Rag2^tm^*^1^*^.1Cgn^*/J; JAX stock #008449) and C57BL/6J were purchased form the Jackson Laboratory. Experiments and procedures were approved by the NIAID Animal Care and Use Committee according to standards set forth in the NIH guidelines, Animal Welfare Act and U.S. Federal law. Euthanasia was carried out using carbon dioxide inhalation in accordance with the American Veterinary Medical Association guidelines (2013 Report of the AVMA panel on euthanasia).

### Vaccines and Challenge Virus

MVA ^26^ was purified by sucrose gradient centrifugation and titrated on primary chicken embryo fibroblasts. 10^7^ PFU of MVA in a volume of 0.5 ml was administered IM. The dose of mRNA-1769 ^8^ was 8 µg, unless stated otherwise, administered IM in a volume of 50 µl. WRvFire ^43^ purified by sucrose gradient sedimentation was inoculated IN in amounts ranging from 10^4^ to 10^6^ PFU in 20 µl of phosphate buffered saline with 0.05% bovine serum into one nostril. For IR, 10^6^ PFU of WRvFire was administered in a volume of 15 µl using a syringe and the animals were held upside down for 2 min to prevent leakage of the material. Percutaneous inoculation of 10^5^ PFU in a volume of 10 µl was administered at the base of the tail by 20 skin scratches with a 25-gauge needle.

### Passive serum transfer

Pooled heat inactivated serum for passive transfer was obtained from cynomolgus macaques that had been primed and boosted with 15 µg, 50 µg, or 150 µg of mRNA-1769. BALB/c, C57Bl/6 and Rag2 KO mice received 0.5 ml of immune or control serum IP one day prior to or one day following IN challenge with 10^5^ PFU WRvFire. The day following transfer, mice were bled to determine levels of VACV neutralizing antibody. Mice were imaged and weighed over the next 2-4 weeks.

### Neutralization and comet inhibition assays

Virus neutralization assays were carried out by incubating sucrose gradient purified VACV WR that expresses GFP with serum dilutions as previously described ^44^. For the comet inhibition assay, BS-C-1 monolayers in 12-well dishes were incubated with diluted samples of VACV strain IHD-J for 1 h at 37°C. The medium was aspirated, and the monolayers washed with fresh medium to remove free virus. The plates were further incubated at 37°C for 48 h and then fixed and stained with crystal violet.

#### BL imaging

An IVIS 200 system (Perkin Elmer, Waltham, MA) was used to image infected animals as described ^45^. Image collection times of up to 60 sec and binning factors were held constant for all animals in an experiment. Photon flux (photons/s/cm2/sr) was determined for regions of interest including the head, body, and areas around rectum and percutaneous infection. Acquisition and analysis were performed with Living Image software (Perkin Elmer).

## ACKNOWLEDGEMENTS

Research was supported by the Division of Intramural Research, NIAID. Animal care was provided by the NIAID Comprehensive Medical Branch. The opinions or assertions contained herein are the private views of the authors, and are not to be construed as official, or as reflecting true views of the Department of the Army, the Department of Defense, or the US Government

## AUTHOR CONTRIBUTIONS

Conception and design of experiments: BM, AWF

Investigation: CAC, MAI, JLA, PLE

Materials: AWF, JH, EMM

Project Administration: BM, AWF, TRF, AC

Supervision: BM, AWF

Original draft and figures: BM, CAC

Comments and editing of manuscript: All authors

